# Transduction catalysis: doxorubicin accelerates and enhances rAAV-mediated gene expression in the cortex of mouse, cat and monkey

**DOI:** 10.1101/2020.06.08.139667

**Authors:** Hongliang Gong, Nini Yuan, Zhiming Shen, Cheng Tang, Stewart Shipp, Liling Qian, Yiliang Lu, Ian Max Andolina, Shenghai Zhang, Jihong Wu, Hui Yang, Wei Wang

## Abstract

Rapid and efficient gene transduction via recombinant adeno-associated viruses (rAAVs) is highly desirable across many basic and clinical research domains. Here we report vector co-infusion with doxorubicin, a clinical anti-cancer drug, markedly enhanced rAAV-mediated gene expression in the cerebral cortex across mammalian species (cat, mouse, and macaque), acting throughout the time-period examined and detectable at just three days post-transfection. This enhancement showed serotype generality, being common to rAAV serotypes 2, 8, 9 and PHP.eB tested, and was observed both locally, and at remote locations consistent with doxorubicin undergoing retrograde axonal transport. All these effects were observed at doses matching human blood plasma levels in clinical therapy, and lacked detectable cytotoxicity as assessed by cell morphology, activity, apoptosis and behavioral testing. Altogether, this study identifies an effective means to improve the capability and scope of *in vivo* rAAV applications, accelerating and augmenting gene transduction at doxorubicin concentrations paralleling medical practice.

**Highlights:**

1. Anti-cancer drug doxorubicin doubles the rate of rAAV-mediated transgene expression
2. Doxorubicin enhancement generalizes across rAAV serotypes and animal species
3. The effect is observed in both locally and retrogradely infected cortical neurons
4. The effective dosage is free from appreciable cytotoxicity and matches clinical settings

## INTRODUCTION

Manipulation of gene expression in specific populations of neurons is a potent capability for both neuroscientific research (Sun and Schaffer, 2018) and therapeutic intervention of genetic disorders (Deverman et al., 2018). Vectors derived from adeno-associated virus (AAV) are especially favored for *in vivo* gene manipulation as they provide a non-pathogenic and minimally immunogenic means to achieve persistent, stable transgene expression within mammalian central nervous system (CNS) (Hudry and Vandenberghe, 2019; Kaplitt et al., 1994). Combined with molecular genetic techniques, recombinant AAV (rAAV) has proved to be a powerful research tool, widely used in gene editing and expression modulation (Ran et al., 2015), neuronal morphology mapping (Parekh and Ascoli, 2013), *in vivo* imaging (Tian et al., 2009), neural circuit analysis (Zhang et al., 2016) and treatment of neurological disorders (Hudry and Vandenberghe, 2019). However, the expression of genes transduced via rAAV undergoes a prolonged lag phase before reaching significant levels for intervention (≥ two weeks) (Diester et al., 2011; Ju et al., 2018). This delay in expression limits the utility of rAAV applications in both experimental and clinical practice, especially in neurodevelopmental studies (Krol and Feng, 2018) and disease treatments that require timely intervention to forestall irreversible tissue damage (e.g., ischemic brain injury) (Sehara et al., 2018). Thus a more effective means to induce rapid and high-level rAAV-based transgene expression in mammalian CNS is in much need of development.

Previous research efforts have aimed to improve the vector system, i.e. the viral capsid and genome. By utilizing natural isolates with different tropisms, new rAAV variants with superior efficacy have been engineered through capsid transformation (Broekman et al., 2006), amino acid mutation (Zhong et al., 2008), domain/subunit swapping (Rabinowitz et al., 2004), or screened from mutant capsid libraries (Deverman et al., 2016; Maheshri et al., 2006; Yang et al., 2009). At the level of vector genome, the vector expression cassette has been optimized with stronger promoters, enhancers, and more effective polyadenylation sequences (Monahan et al., 2015; Peel et al., 1997). By modifying one of the inverted terminal repeats (ITR), self-complementary AAV (scAAV) has been generated to bypass second-strand synthesis to improve the efficacy of transduction (McCarty et al., 2003). Combined with advances in delivery techniques, good progress has been achieved in respect of transduction efficiency in small animals (Chan et al., 2017). However, for large mammals represented by non-human primates and humans, it remains challenging to achieve consistent, high levels of transduction across volumetrically extensive brain structures. This problem is manifest in both therapeutic trials and research applications (Hadaczek et al., 2010; Hwu et al., 2012), enforced by capsid specific cytotoxic T lymphocyte response (Mingozzi et al., 2007). Furthermore the delay in expression remains critically important, since scAAV can only be used for smaller transgenes (less than 2.2 kb) (Wang et al., 2019).

The application of exogenous agents is a new strategy to promote rAAV transduction in the CNS. Early studies revealed that techniques such as adenovirus co-infection, ionizing radiation and genotoxic agents can all promote rAAV2 transduction, inducing faster and higher gene expression in cultured cells (Alexander et al., 1994; Fisher et al., 1996; Kanazawa et al., 2001). Similar promotion was also achieved with chemical agents that damage DNA, inhibit topoisomerase activity, or modulate proteasome function (Nicolson et al., 2016; Russell et al., 1995; Yan et al., 2004). These agents appear to promote viral intracellular trafficking and/or second-strand DNA synthesis, two rate-limiting steps in rAAV transduction that see little improvement with vector capsid engineering. However, severe side-effects have hindered their application *in vivo*, calling for alternative drugs or strategies that are less cytotoxic. ***Doxorubicin*** is a frontline drug that has been widely used in cancer therapy (Tacar et al., 2013). As a known inhibitor of topoisomerase and proteasome (Sishi et al., 2013; Tacar et al., 2013), it has clear potential to promote rAAV transduction, as evidenced by application to cultured cells (Yan et al., 2004; Zhang et al., 2012; Zhang et al., 2009). However, to pave the way for incorporation of doxorubicin in rAAV protocols for CNS, it is essential to examine the precise time-course of neural transduction, and the dose-dependency of the long-term balance between efficacy and neurotoxicity. It was thus our goal to determine just how effectively doxorubicin might accelerate rAAV-mediated gene transduction in mammalian CNS and establish the ceiling for toxicity. Furthermore, it is imperative to determine the universality of the action of doxorubicin across a range rAAV serotypes, brain structures, and mammalian species.

Here, we have systematically evaluated the effect of doxorubicin on transgene expression in the cerebral cortex of mice, cats, and macaque monkeys, by intraparenchymal administration of doxorubicin in combination with different rAAV serotypes (rAAV8, rAAV2, rAAV9 and rAAV-PHP.eB). We found that doxorubicin greatly enhanced transgene expression mediated by every rAAV vector that we tested, differing little between species. Across subjects, doxorubicin took almost immediate effect and continued to act throughout the time period examined. Dose escalation testing showed that the effect of doxorubicin on rAAV transduction was concentration-dependent, and was mediated with negligible adverse side-effects at doses well below tolerance. Finally, by co-administration with rAAV2-retro for tracing connections in macaque visual cortex, we found that doxorubicin could also enhance rAAV transduction at remote locations, consistent with retrograde axonal transport from the site of injection. In sum, our findings offer a feasible strategy that can complement vector system optimization for rapid and efficient *in vivo* rAAV transduction in the mammalian CNS.

## RESULTS

### Doxorubicin Enhances rAAV Transduction with No Impact upon Cell Tropism

We began by evaluating the effect of doxorubicin in the cerebral cortex of cat, a well-studied mammalian model (Vite et al., 2003). For our standard vector we selected rAAV8, a serotype isolated from rhesus monkey with the express aim of developing a vector for gene therapy with minimal immunogenicity in human tissues (Gao et al., 2002). rAAV8 proved to have a CNS transduction capability superior to that of rAAV2 (Broekman et al., 2006) and could mediate efficient transduction after intraparenchymal injection in various cortical structures of primate brains (Gilkes et al., 2016; Masamizu et al., 2010). rAAV8-hSyn-EGFP, a rAAV8 vector which encoded green fluorescent protein (GFP) under the neuron-specific promoter hSyn, was constructed and infused into cat visual cortex via stereotactic injection at a titer of 1×10^12^ vector genome per milliliter (vg/ml, 1 μl per injection), combined with 10 μg/ml doxorubicin (left hemisphere) or vector alone (right hemisphere) (**Figure 1A**). GFP expression was determined with standard histological techniques thirty days later. As revealed by fluorescence images of brain sections, pressure-injection of vector solution led to focal transduction within a radius of 750 μm, with GFP expression in cell bodies and axons projecting to extrinsic structures (**Figure 1B)**. Compared with the control group, GFP fluorescence in doxorubicin-treated group was noticeably brighter, with intensity significantly increased (∼2.5-fold) by doxorubicin (1015 ± 58.4 vs 405.5 ± 46.1, p < 0.001; **Figure 1C**). Consistent with studies in mouse and marmoset (Watakabe et al., 2015), rAAV8-mediated GFP expression was not homogeneous across cortical layers, with layer 4 the weakest. As indicated by neuronal-specific marker NeuN, confocal images revealed that the proportion of GFP-expressing neurons was markedly increased, exhibiting higher fluorescence across all cortical layers when treated with doxorubicin (layer 2/3: 92.5 ± 1.5% vs 81.7 ± 2.1%; layer 4: 79.4 ± 2.8% vs 41.3 ± 2.8%; layer 5: 92.2 ± 2.2% vs 78.1 ± 1.7%; layer 6: 91.6 ± 1.6% vs 75.4 ± 2.5%, all p < 0.01; **Figure 1D**). Across all layers the average ratio of GFP-positive neurons was 90 ± 2% in doxorubicin-treated group, compared with 70 ± 2% in the control group (**Figure 1D**). More notably, after a tenfold decrease of vector titer at 1×10^11^ vg/ml, doxorubicin treatment still gave an equivalent level of GFP expression to the control at 1×10^12^ vg/ml (403.3 ± 32.8 vs 405.5 ± 46.1).

**Figure 1.**
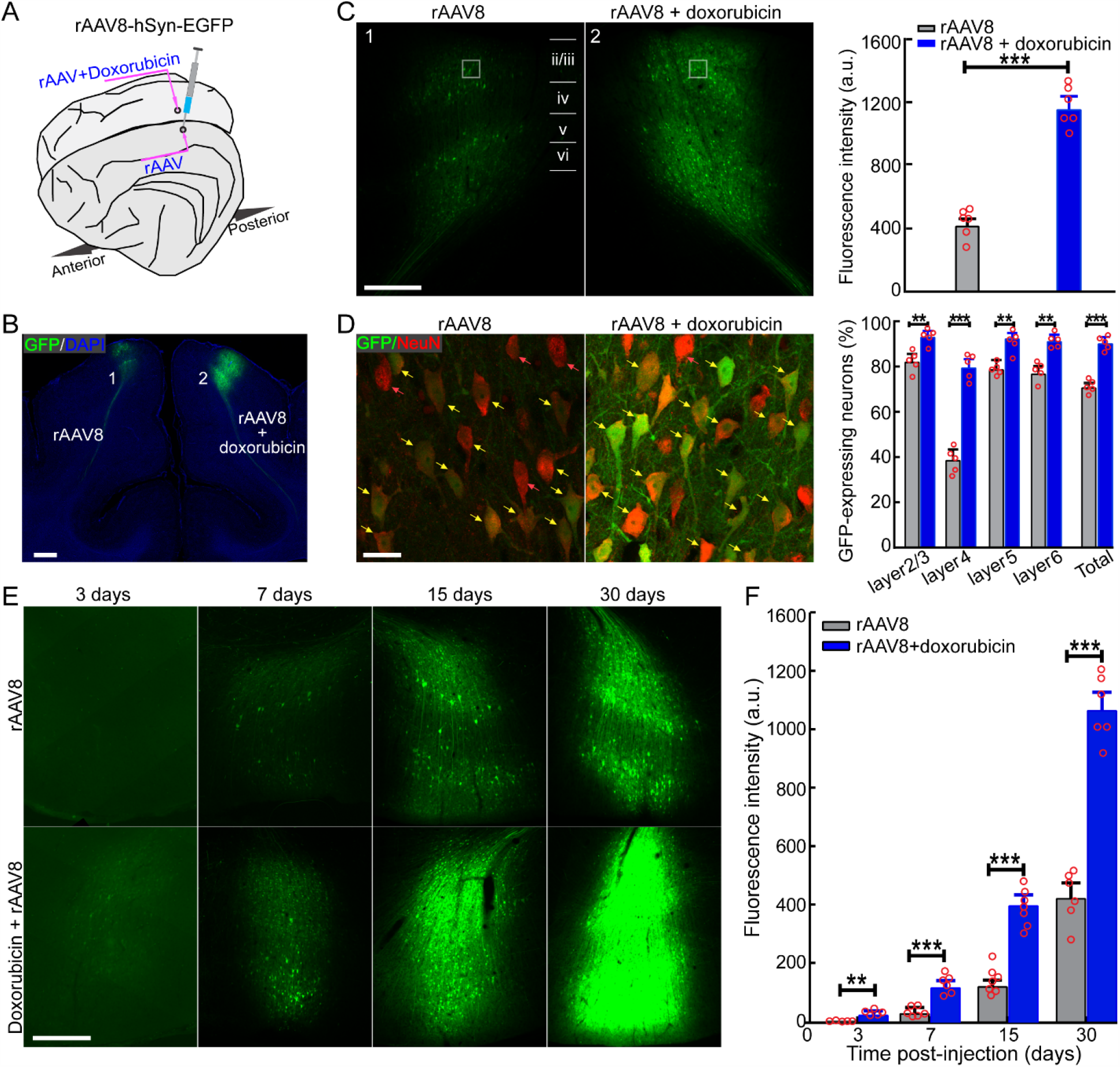
Doxorubicin Promotes and Accelerates rAAV8-Mediated Gene Transduction in Cat Visual Cortex. (**A**) Schematic representation of injections. rAAV8-hSyn-EGFP alone or coupled with doxorubicin were delivered into the left and right hemisphere of cat visual cortex, and GFP fluorescence was examined after a period of expression. (**B**) Representative image of GFP fluorescence in the control and doxorubicin-treated sites. Blue, DAPI; green, GFP. Scale bar, 1 mm. (**C**) Cortical fluorescence intensity comparison after 30 days of expression. 6 injections from 3 cats per group; unpaired two-tailed t-test. Scale bar, 500 μm. (**D**) Transduction efficiency assessed by colocalization of GFP and NeuN marker. Left: confocal images of GFP expression and NeuN staining of layer 2/3 (boxed in **C**). Yellow arrows, GFP-positive neurons; red arrows, GFP-negative neurons; Scale bar, 25 μm. Right: quantification of transduction efficiency across cortical layers. 5 injections from 3 cats per group; unpaired two-tailed t-test. (**E**) Representative image of GFP expression at the specified time points. The gamma value was adjusted equally to enable the visualization of GFP signals at 3 days. Scale bar, 500 μm. (**F**) Quantification of GFP intensity at varied intervals post-injection. 5-6 injections per group; unpaired two-tailed t-test; data are presented as mean ± SEM. *p <0.05, **p <0.01, ***p <0.001.

To determine whether the effect of doxorubicin on rAAV transduction is exclusive to neurons, we replaced hSyn with an ubiquitous promoter CAG. The modified viral vector, rAAV8-CAG-EGFP (titer: 5×10^11^ vg/ml) was administered alone or combined with 10 μg/ml doxorubicin. The phenotypes of transduced cells were determined by immunohistochemical detection of cell-type-specific markers seven days later, i.e. NeuN for neurons, glial fibrillary acidic protein (GFAP) for astrocytes and ionized calcium binding adaptor molecule (Iba-1) for microglia (**Figure S1A-C**). As anticipated, we found that under the control of CAG promoter GFP was expressed not only in neurons (NeuN^+^) but also in astrocytes (GFAP^+^); fluorescence intensity (**Figure S1D**) and the proportions of both GFP-positive neurons and astrocytes were significantly increased at the injection sites of doxorubicin-treated group relative to the control (**Figure S1E**). With or without doxorubicin, there was no detectable fluorescence signal in microglia (**Figure S1C** and **S1E**), consistent with experience to date that this cell class is relatively refractory to transduction by rAAV vector (Maes et al., 2019). These results suggest that doxorubicin treatment enhances neuronal transduction mediate by rAAV8 vector, with no quantitative impact upon its cellular tropism, which is governed by vector’s capsid (Srivastava, 2016).

### Doxorubicin Accelerates rAAV-Mediated Gene Expression

Practical, rapid onset of rAAV-mediated gene expression is highly significant in regard to studies of brain development, as well as treatment of diseases that require timely intervention (Krol and Feng, 2018; Shimazaki et al., 2000). To explore the impact of doxorubicin on the onset of rAAV transduction *in vivo*, we carefully examined the time course of transgene expression in doxorubicin-treated versus control groups. Following our established procedure, rAAV8-hSyn-EGFP alone or coupled with 10 μg/ml doxorubicin was infused into cat visual cortex, and the expression levels of GFP were determined with standard histological techniques at 3, 7, 15 and 30 days following injection (**Figure 1E**). We found that the pace of transgene expression was greatly accelerated in the presence of doxorubicin. In the control group, GFP fluorescence could be detected no earlier than 7 days post-injection. This reduced to 3 days for doxorubicin-treated group, with a steady increase in GFP expression thereafter. At 7 days after injection, GFP activity in doxorubicin-treated group was about 4-fold higher than that in the control group (104.3 ± 9.6 vs 25.2 ± 2.5, p < 0.001), actually matching the 15-day level of the control group (104.3 ± 9.6 vs 118.7 ± 10.9)(**Figure 1E**). Throughout the period of examination, the level of GFP expression in doxorubicin-treated group consistently exceeded the control group (3 days: p = 0.0015; 7 days: p = 0.0003; 15 days: p < 0.0001; 30 days: p < 0.0001; **Figure 1F**). These results reveal that doxorubicin significantly accelerates rAAV transduction, facilitating timely interventions in the CNS by means of rapid and efficient transgene expression.

### Dose-Dependency for Doxorubicin Enhancement of rAAV-Mediated Gene Expression

To determine the optimal dosage of doxorubicin for *in vivo* neurological administration, a trade-off between efficacy for rAAV transduction versus toxicity to neural tissues, rAAV8-hSyn-EGFP was infused into cat cerebral cortices at a titer of 1×10^12^ vg/ml in combination with varied concentrations of doxorubicin (range 0-100 μg/ml). GFP expression was examined with fluorescence microscopy seven days post-injection. As shown in **Figure 2A**, GFP expression levels were closely tied to doxorubicin concentration, displaying a steady increase over a range of 0.1 to 30 μg/ml. To mitigate doxorubicin concentration loss through diffusion, GFP intensity was measured within a ROI of 500 × 1000 μm at the core of each injection, as indicated in the control (**Figure 2A**). Quantitative analysis confirmed the dose-dependent enhancement of rAAV-mediated GFP expression by doxorubicin, with the maximum obtained at 30 μg/ml (Control vs 1.0 μg/ml, p = 0.02; control vs 3.0 μg/ml, p = 0.008; control vs 10 μg/ml, p = 0.002; control vs 30 μg/ml, p = 0.0012; **Figure 2B**). For the topmost doxorubicin dosage of 100 μg/ml, GFP intensity decreased sharply at the core of injection site, demonstrating notable cytotoxicity.

**Figure 2.**
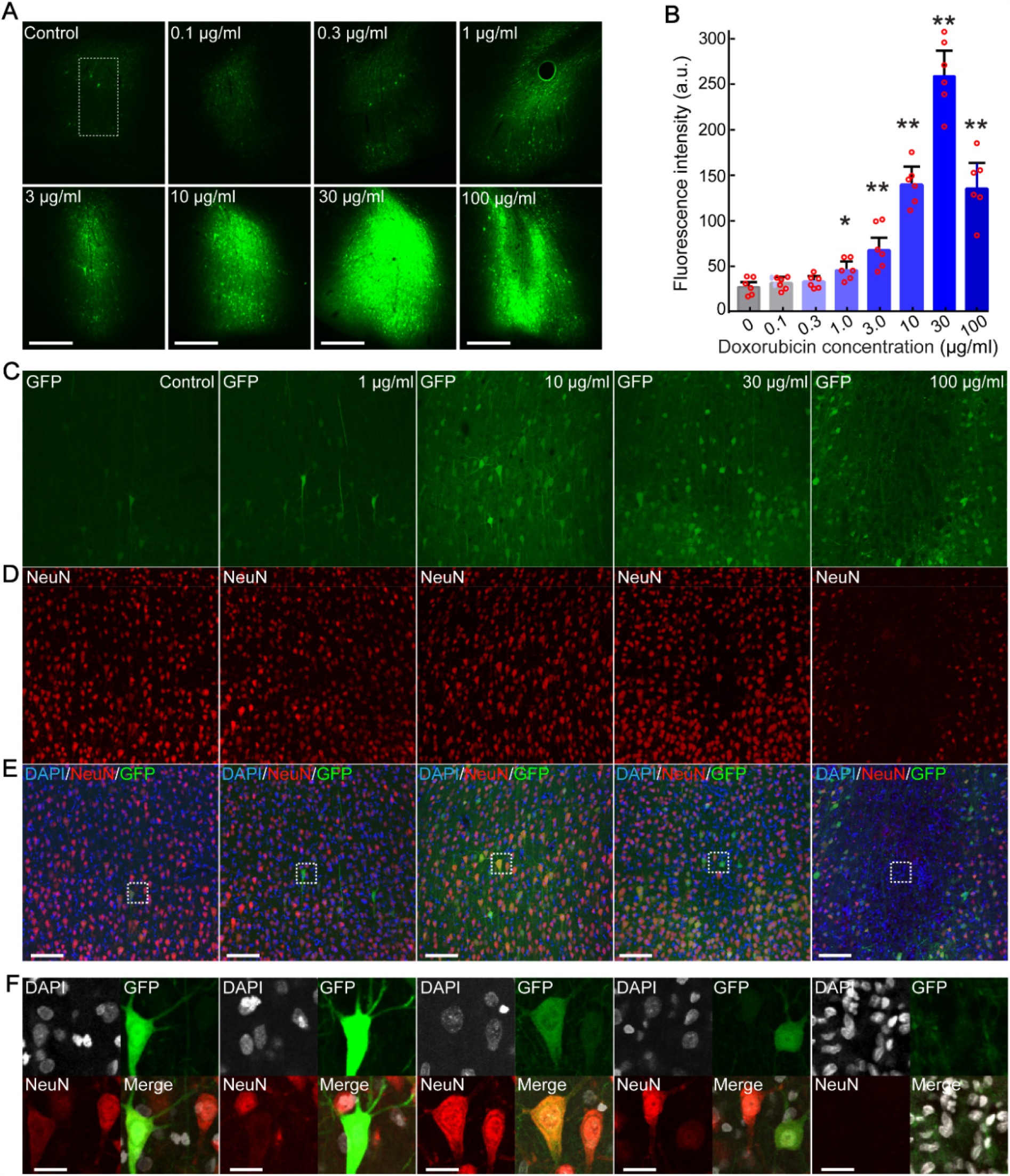
The Effect of Doxorubicin on rAAV Transduction Is Concentration-Dependent, and Free from Appreciable Cytotoxic Effects at a Dosage from 1 to 10 μg/ml. (**A**) Representative fluorescence images of injections seven days after co-infusion of rAAV8-hSyn-EGFP with varied concentrations of doxorubicin. Scale bar, 500 μm. (**B**) Quantification of core fluorescence intensity (as indicated in the control of **A**). 6 injections from 6 cats per group. One-way ANOVA, post hoc Tukey Kramer correction; mean ± SEM, *p < 0.05, **p < 0.01. (**C-E**) Status of transduced neurons (GFP, green) at cellular level (**C**), quantified by neural marker NeuN (**D**), and nuclear marker DAPI (**E**). Scale bar, 100 μm. (**F**) Magnified views of neuronal cells under different doxorubicin concentrations (as indicated in the inset of **E**). Scale bar, 20 μm.

To further substantiate the differential dose-effect of doxorubicin, we analyzed the status of GFP-expressing neurons with the nuclear marker DAPI and the neuron-specific marker NeuN (**Figure 2C-F**). Confocal images revealed that cell bodies and neurites of infected neurons exhibited intact and normal morphology after doxorubicin treatment at a dosage from 0.1 to 10 μg/ml. At a dose of 30 μg/ml the neurites of neurons in the core of injections receded, and cell bodies became round with weak or even absent NeuN signals. When doxorubicin concentration reached 100 μg/ml neurons were completely absent from the core of the injection, with glial accumulations in their place. These data further informed the optimal dosage for use of doxorubicin in the CNS, indicating a ceiling concentration of 10 μg/ml, up to which rAAV-mediated gene transduction is notably enhanced by doxorubicin with negligible cytotoxicity to neural tissues.

### Locally Injected Doxorubicin Lacks Appreciable Cytotoxicity for Neural Tissue

To establish the safety of *in vivo* neurological administration of doxorubicin at 10 μg/ml, we set out to determine whether or not this concentration has apparent cytotoxic effects upon neural tissues. First, we used TUNEL assay to assess neuronal apoptosis consequent to doxorubicin application. Seven days after delivery of doxorubicin and rAAV vector, TUNEL staining was performed on brain sections and apoptotic cells were screened by confocal fluorescence microscopy. As presented in **Figure 3A**, no appreciable apoptotic effect was observed in either doxorubicin-treated group or the control group (rAAV alone), when compared to the positive control treated with DNase I, as indicated by fluorescent-labeled nuclear marker DAPI.

**Figure 3.**
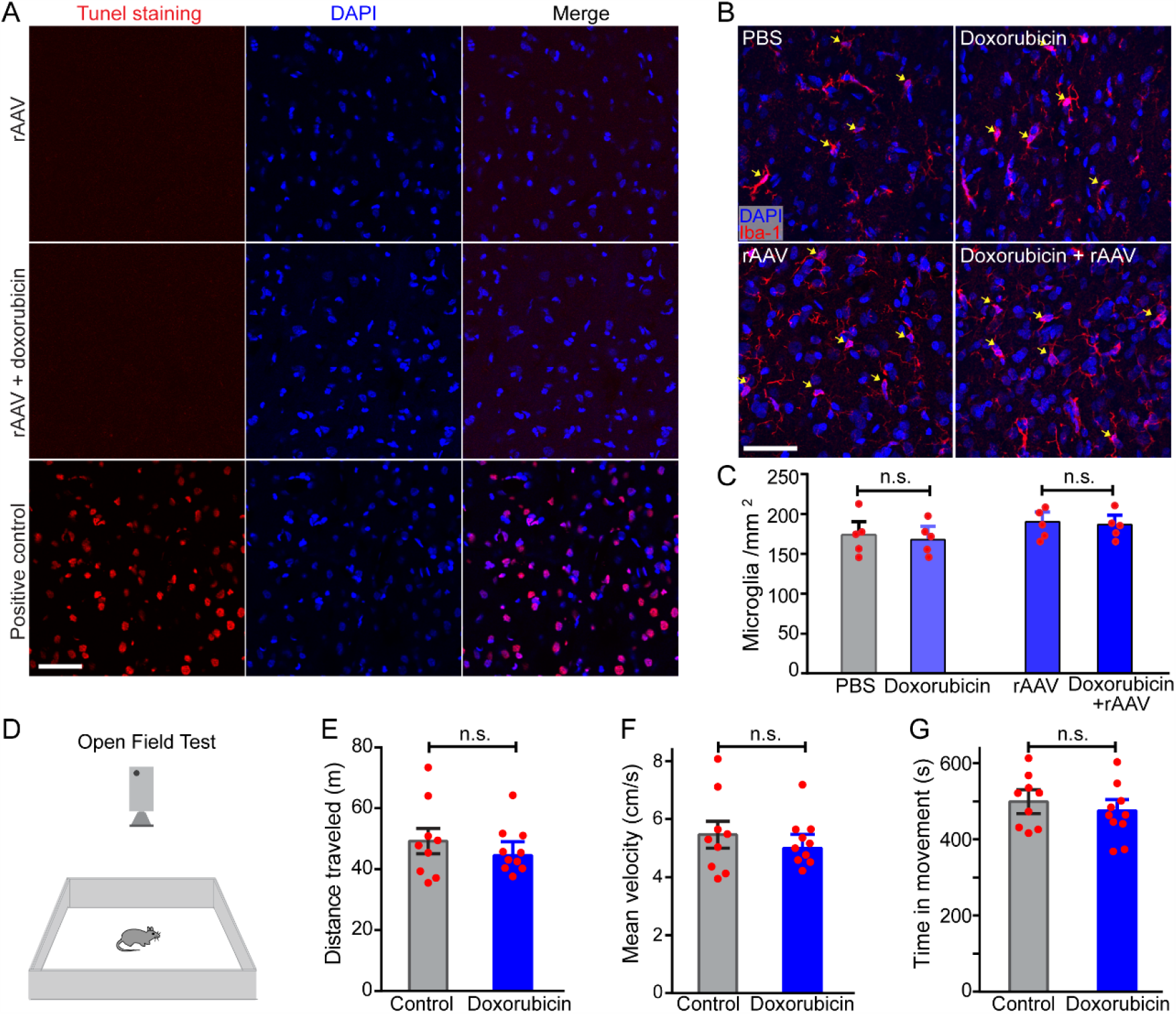
Doxorubicin Induces No Appreciable Side Effects on Neural Tissue under Experimental Conditions. (**A**) Confocal images of TUNEL stained sections seven days after administration of rAAV8 alone or with doxorubicin. Positive control samples pretreated with DNase I. Blue, DAPI; red, apoptotic cell. Scale bar, 50 μm. (**B**) Representative images of fluorescent-labeled microglia cells (yellow arrows) by immunohistochemical detection of Iba-1; blue: DAPI, red: Iba-1. Scale bar, 50 μm. (**C**) Quantification of microglia cell numbers in the test sites. 5 injections per group. Unpaired two-tailed t-test; mean ± SEM. (**D**) Schematic diagram of open field test. rAAV8-hSyn-EGFP vector was administered into mouse M1 with or without doxorubicin, and mice were placed in an open field chamber (40 × 40 cm) to allow 15 minutes for free exploration 15 days post-injection. (**E-G**) Quantification of distance traveled (Wilcoxon rank-sum test), mean velocity (Wilcoxon rank-sum test) and time in movement (Unpaired two-tailed t-test) between doxorubicin-treated mice and the control. For quantification: n = 9 and n =10 mice in the control and doxorubicin groups, mean ± SEM, n.s. p > 0.05.

In addition to apoptosis, doxorubicin therapy can also cause inflammation, a major contributor to its cardiotoxicity and other side-effects (Sauter et al., 2011). To further establish the feasibility of doxorubicin treatment, the inflammatory response was examined by immunohistochemical detection of microglia, which generally aggregates in inflamed and necrotic tissues (Vela et al., 2002). After a similar interval, i.e. seven days following injection of rAAV and doxorubicin, microglia were detected and counted in brain sections by antibody recognition of its specific marker Iba-1 (**Figure 3B**). Quantitatively, the outcome revealed no difference between the doxorubicin and control groups in regard to the density of microglia after a single injection (164.0 ± 7.5/mm^2^ vs 173.2 ± 11.8/mm^2^, p = 0.53; **Figure 3C**). Co-administration with rAAV8 vector gave a similar result, i.e. the rAAV-plus-doxorubicin group versus rAAV group (186.2 ± 6.2/mm^2^ vs 189.6 ± 5.9/mm^2^, p = 0.7; **Figure 3C**), indicating that there was no inflammatory response following a single injection of doxorubicin.

Finally, to determine whether doxorubicin treatment might induce any side effects at the behavioral level, we co-infused rAAVs with doxorubicin or vector alone (as a control) into the primary motor cortex (M1) of mice in order to determine any consequent change in locomotor activity. After a period of fifteen days, the mice were placed in an open field chamber and allowed 15 minutes for free exploration, monitored by video camera (**Figure 3D**). None of the locomotor parameters analyzed revealed any significant difference in behavior between doxorubicin-treated mice and the control: specifically, total distance traveled (4691 ± 1332 vs 4295 ± 736 cm, p = 0.28; **Figure 3E**), mean velocity (5.2 ± 1.5 vs 4.8 ± 0.8cm/s, p = 0.28, **Figure 3F**) and time in movement (8.28 ± 0.37 vs 7.77 ± 0.38 min, p = 0.34; **Figure 3G**) were all equivalent. Taken together, these data show that local injection of doxorubicin lacks detectable adverse effects upon neural tissue under typical experimental conditions.

### Doxorubicin Enhances rAAV Transduction Irrespective of rAAV Serotypes and Animal Species

AAV has multiple serotypes and new variants of rAAV are continually being developed for experimental and clinical purposes (Wang et al., 2019). To investigate how far our findings represent a general agonist effect of doxorubicin upon rAAV transduction, we employed two other rAAV serotypes: rAAV2 and rAAV-PHP.eB, the latter a variant of AAV9 selected to maximize CNS transduction via intravenous administration (Chan et al., 2017). Combined with 10 μg/ml doxorubicin, viral vectors of rAAV2, rAAV8 and rAAV-PHP.eB encoding GFP under the CAG promoter were injected into cat cerebral cortex and GFP expression was assessed seven days later. We observed similar levels of GFP expression in the control groups of rAAV8 and rAAV-PHP.eB, each significantly better than that achieved by rAAV2 (**Figure 4A**). More importantly, GFP intensity (rAAV2, p = 0.006; rAAV8, p = 0.0094; rAAV-PHP.eB, p = 0.012; **Figure 4B**) and percentage of transduced cells (rAAV2: 15 ± 1% vs 5 ± 1%, p = 0.0086; rAAV8: 37 ± 4% vs 16 ± 1%, p = 0.0094; rAAV-PHP.eB: 35 ± 3% vs 13 ± 1%, p = 0.0005; **Figure 4C**) were both significantly upregulated by doxorubicin in all three serotypes of rAAV, suggesting doxorubicin indeed has a general agonist effect upon gene transduction mediated by rAAV vectors, irrespective of vector serotype.

**Figure 4.**
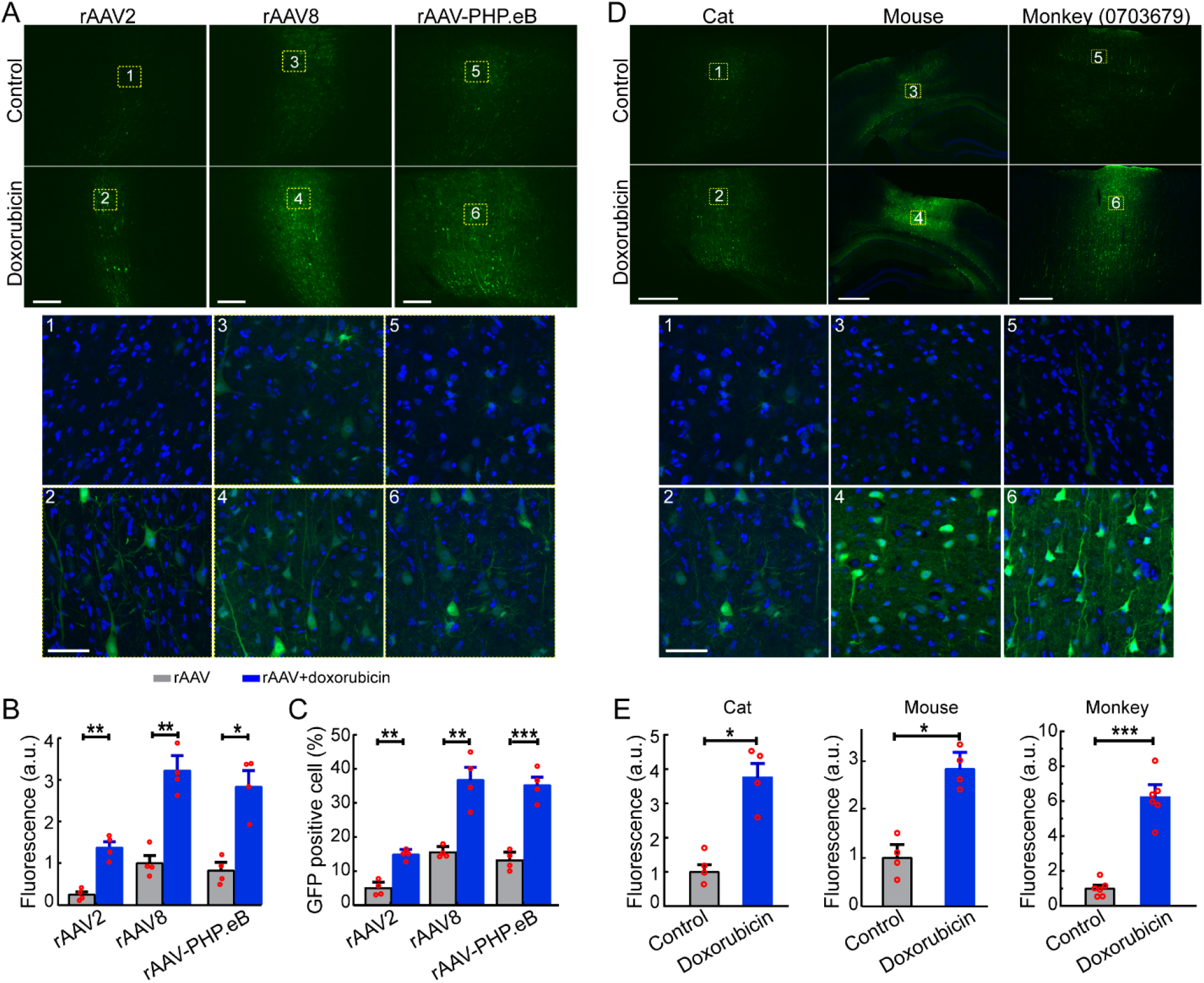
The Agonist Effect of Doxorubicin Is Reproducible with Different rAAV Serotypes and Animal Species. (**A**) Representative images showing GFP expression via viral vectors of rAAV2, rAAV8 and rAAV-PHP.eB under the CAG promoter. Upper panel: images of the injection sites pertaining to rAAV2, rAAV8 and rAAV-PHP.eB vector, respectively. Scale bar, 250 μm. Lower panel: magnified views of the boxed regions 1-6. DAPI: blue, GFP: green. Scale bar, 50 μm. (**B,C**) Quantification of mean GFP intensity and percentage of transduced cell. 4 injections from 4 cats per group; unpaired two-tailed t-test. (**D**) Representative images of vector injection-sites from the cerebral cortex of cat, mouse and macaque monkey. rAAV-PHP.eB-CAG-EGFP vector was used and GFP expression was assessed after seven days of expression. Lower panel: magnified views of the boxed regions 1-6. DAPI: blue, GFP: green. Scale bar, 500 μm (upper), 50 μm (lower). (**E**) Quantification of mean GFP intensity in cat, mouse and monkey. For quantification, 4 injections from 4 animals per group in cat and mice, and 6 injections per group from two monkeys; unpaired two-tailed t-test; mean ± SEM, *p <0.05,**p <0.01. ***p <0.001.

We further tested the effect of doxorubicin in the cerebral cortex of different animal species. After an infusion of rAAV-PHP.eB vector into the cerebral cortex of mouse (visual cortex), cat (visual cortex) and macaque monkey (motor cortex)(**Figure 4D**), we obtained a substantial, quantifiable enhancement of transgene expression across all three mammalian species under doxorubicin treatment (Cat, p = 0.012; Mouse, p = 0.011; Monkey, p = 0.0006; **Figure 4E)**. Qualitative examples of the enhancement of doxorubicin in macaque motor cortex are further shown in **Figure S2**. These results indicated that doxorubicin may well exert a general effect upon rAAV transduction, regardless of rAAV serotype and host species.

### Doxorubicin Enables Rapid Gene Transduction for *In Vivo* Two-Photon Imaging

Two-photon imaging has been a powerful tool widely used across many fields including: physiology (Cahalan and Parker, 2008), neurobiology (Svoboda and Yasuda, 2006) and tissue engineering (Rubart, 2004). Exploiting rAAV vectors carrying genetically encoded indicators, two-photon Ca^2+^ imaging has lent insight into the structural dynamics and functional activity of neuronal populations at single-cell level (Seidemann et al., 2016). However, as rAAV transduction ensues, Ca^2+^ fluorescence signal strength is rarely adequate to image cortical activity earlier than two weeks post-injection (Attinger et al., 2017; Iacaruso et al., 2017). Based on our anatomical findings we predicted that doxorubicin treatment should reduce this delay period.

To examine this, we constructed rAAV9-hSyn-GCaMP6s vector (titer: 2×10^12^ vg/ml, 1 μl per site) and infused it into mouse primary visual cortex (V1) in combination with doxorubicin or alone. As shown by **Figure 5A** for a large window of brain surface, the fluorescence signal in doxorubicin-treated sites was evident at the sixth day after injection, at which time point it was undetectable in the control site. Quantitative measurements of fluorescence signal revealed that doxorubicin more than doubled the rate of expression of GCaMP6s, reaching a plateau at 30 days, whereas the control mice achieved 50% of this level of expression much later, at around 60 days (6 day, p = 0.003; 15 day, p = 0.0133; 30 day, p = 0.0003; 60 day, p = 0.0011; **Figure 5B**). When examined with two-photon microscopy at the sixth day, the morphology of cell bodies was clearly observable under doxorubicin-treated conditions. Accordingly, using drifting sine-wave gratings as visual stimuli, robust direction-selective responses of mouse V1 neurons could be recorded in doxorubicin-treated areas, but not in the control (**Figure 5C,D**); comparable neural activities in untreated mice could only be obtained until two weeks post-administration. Meanwhile, at a comparative expression level of GCaMP6s, e.g. the 15th day of doxorubicin-treated versus the 30th day of control sites (**Figure S3A,B**). We found no significant difference between two groups in the response strength (**Figure S3C**), with only a slight reduction in the orientation selectivity of V1 neurons (**Figure S3D**). These results imply that doxorubicin treatment results in no observable impairment of neural function.

**Figure 5.**
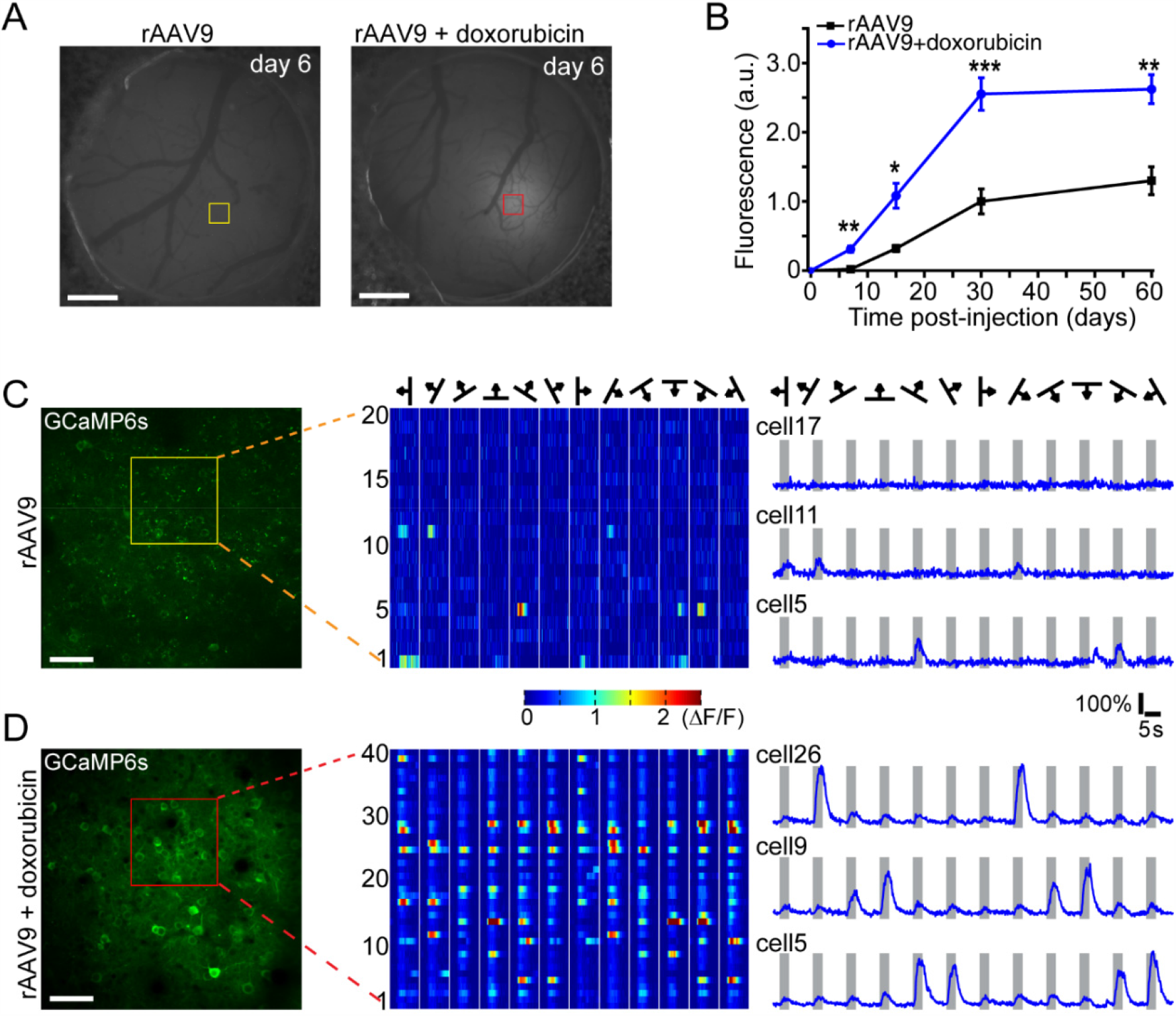
Doxorubicin Induces a Rapid Onset of Transgene Expression for *In Vivo* Study of Neural Function. (**A**) Representative fluorescence images of mouse V1 six days after vector administration (rAAV9-hSyn-GCaMP6s vector: 2×10^12^vg/ml, 1 μl per injection). Scale bar, 1 mm. (**B**) *In vivo* tracking of GCaMP6s expression following administration in mouse visual cortex. 5 mice per group; unpaired two-tailed t-test, mean ± SEM. *p < 0.05. **p < 0.01. ***p < 0.001. (**C,D**) *In vivo* two-photon image of Ca^2+^ fluorescence assessed after only 6 days of expression at a focal plane of injection sites ∼120 μm below cortical surface, as indicated in **A**. Mean fluorescence changes (ΔF/F) of sampled cells (boxed in **C,D**) evoked by drifting sine-wave grating stimuli. Scale bar, 50 μm.

Together these results demonstrated that doxorubicin not only accelerates transgene expression, enabling earlier commencement *in vivo* two-photon imaging, but also has little effect on cortical neuron functionality in the conditions we tested.

### Doxorubicin Enables Efficient Retrograde Tracing of Connections in Macaque Cortex and Thalamus

Finally, motivated by previous reports that doxorubicin undergoes retrograde transport from uptake at axonal terminals (Bigotte and Olsson, 1982; Koda and Van der Kooy, 1983), we considered that doxorubicin have good prospects for remote circuit-selective enhancement of rAAV transduction and therefore could be used for neural circuit interrogation. For this aim we coupled doxorubicin with rAAV2-retro, a vector recently developed for the express purpose of maximizing retrograde access to projection neurons (Tervo et al., 2016).

We conducted this experiment in the primate, in view of the growing importance of systematic, quantified connectional databases in understanding human brain function (Oligschlager et al., 2019; Wang and Kennedy, 2016). rAAV2-retro-CAG-tdTomato (4×10^13^ vg/ml, 1μl) was delivered bilaterally into area V1, either in combination with doxorubicin (right hemisphere) or vector alone as a control (left hemisphere) (**Figure 6A**). Six weeks after injection, we examined tdTomato expression amongst retrogradely labeled cells in the lateral geniculate nucleus (LGN), as well as cortical areas including V2, V4 and V5/MT that send long-range projections to V1. Before examination, we first confirmed good rAAV2-retro transduction across cortical layers at the injection site of V1 (**Figure 6B**). In the LGN, the principal subcortical source of input to V1, we observed dense clusters of neurons labeled with tdTomato (**Figure 6C**). Consistent with previous reports based on tracer dyes (Angelucci and Sainsbury, 2006; Kennedy and Bullier, 1985), the cell bodies of labeled neurons were concentrated ina narrow strip across the six-layer structure of LGN, surrounded by nerve fibers. Compared with the control side, the number of cells labeled with tdTomato on the right side of LGN was far greater, and tdTomato expression was stronger in both neuron cell bodies and fibers (**Figures 6C, 6D** and **S4A**). Amongst prestriate cortical areas, we found large numbers of tdTomato-labeled cells in V2, concentrated in a retinotopically matching subregion close to the injection site in V1(**Figure 6E**), consistent with previous studies (Perkel et al., 1986; Rockland and Pandya, 1979; Rockland and Virga, 1989; Weller and Kaas, 1983). Compared with the left control side, the number of tdTomato-positive cells in the right V2 was greater, and the fluorescent signal was stronger across serial sections (**Figures 6F** and **S4B**), confirming the enhancement effect of doxorubicin to be mediated via retrograde transport in corticocortical as well as thalamocortical projections. This enhancement became more clear in weak or moderate long-range projections - we detected numerous tdTomato-positive neurons in V4, V5/MT and other visual cortices in the right hemisphere, with little or even no tdTomato signal in the left control side; in this respect, performance of rAAV2-retro unaided by doxorubicin was notably poor in comparison to conventional retrograde tracers (**Figures 6G-I** and **S4C,D**). These results indicate that doxorubicin can induce remote enhancement of rAAV transduction via axonal retrograde transport in both thalamocortical and corticocortical projections, and could thus be a useful tool for analysis of neural circuits in large mammals like the macaque.

**Figure 6.**
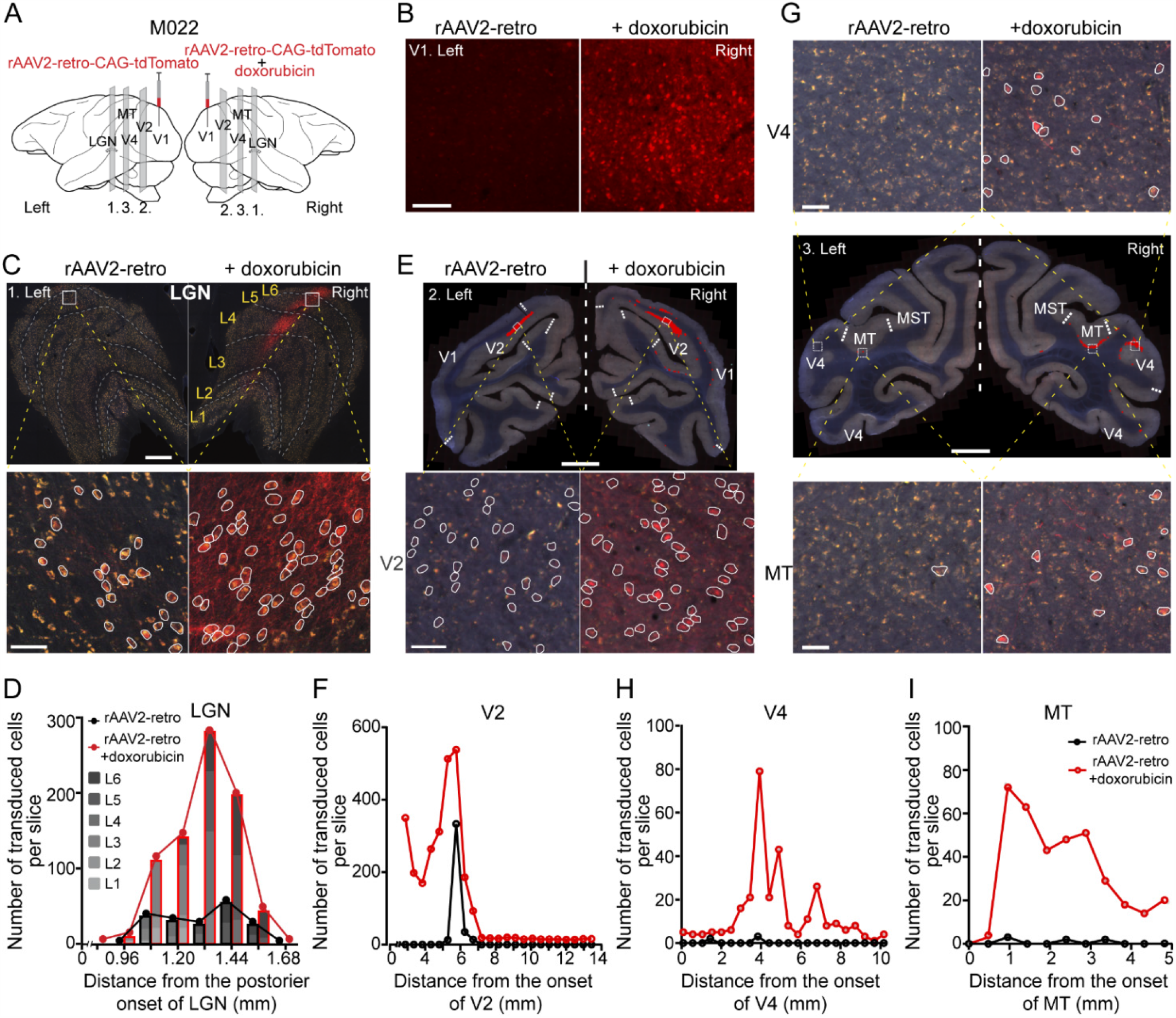
Enhancement of rAAV Transduction via Retrograde Axonal Transport in Macaque Thalamo- and Cortico-Cortical Circuits, See also in Figure S5. (**A**) Schematic showing injections and quantified sources of afferents to V1, including LGN (panel 1), V2 (panel 2), V4 and MT (panel 3). rAAV2-retro-CAG-tdTomato, vector alone or coupled with doxorubicin was infused at symmetrical locations in the left and right hemisphere of monkey V1. Transduced neurons were assessed six weeks later. (**B**) Representative images of neuronal transduction at the injection site. Scale bar, 100 μm. (**C**) Representative images of neuronal transduction in LGN. Transduced neurons from ROI (boxed in the upper panel) were marked by white circles. Scale bar, 500 μm (upper), 50 μm (lower). (**D**) Comparison of retrograde transport efficiency for LGN. Transduced cells were counted in sample sections taken every 120 μm across LGN. (**E**) Representative images of neuronal transduction in V2. Upper panel: whole coronal image (transduced cells are marked with red dots); Lower panel: transduced neurons from ROI (boxed in the upper panel) are marked by white circles. Scale bar, 5000 μm (upper), 50 μm (lower). (**F**) Comparison of retrograde transport efficiency for V2. (**G**) Representative images of neuronal transduction in V4 and MT. Middle panel: whole coronal image of macaque brain (transduced cells are marked with red dots); Top and bottom panels: transduced neurons from ROIs of V4 and MT are marked by white circles. Scale bar, 5000 μm (middle), 50 μm (top and bottom). (**H-I**) Comparison of retrograde transport efficiency for V4 and MT. In **F,H** and **I**, transduced cells were counted in regions of V2, V4 and MT within sample sections taken every 480 μm across the brain from posterior to anterior.

## DISCUSSION

With recent advances in vector engineering, rAAVs have met with great success as carriers for *in vivo* gene transfer (Wang et al., 2019). Clinical trials of rAAV-based gene therapy have demonstrated clear therapeutic efficacy (Christine et al., 2019; Mendell et al., 2017; Mittermeyer et al., 2012). rAAV has also become a powerful tool for cell type-specific gene modulation in many fields of basic neuroscience (Betley and Sternson, 2011). However, it is also recognized that rAAV still has some shortfalls, which include insufficient transduction in targeted tissues and tardiness of gene expression after transduction (Wang et al., 2019). Therefore, more efficient strategies for rapid-onset and high-level transgene expression at lower doses are much to be desired for *in vivo* rAAV transduction. Here we outline how the clinical pharmacological agent ***doxorubicin*** might be exploited to enhance the efficiency of transgene expression in a range of basic and therapeutic research applications across various combinations of rAAV serotype and experimental species.

### Mechanism and Time Course of Doxorubicin Action

After infection of the host cell the rAAV genome has to undergo several processes before transgene expression can occur, amongst which intracellular trafficking and second-strand DNA synthesis are two rate-limiting steps (Ferrari et al., 1996; Harbison et al., 2008). The action of doxorubicin – an inhibitor of topoisomerase II and DNA synthesis (Tacar et al., 2013) – to facilitate this process is therefore paradoxical. A clue may exist in the long-known finding that agents causing DNA damage, including ionizing radiation, have the same general capability as topoisomerase-inhibitors to enhance the efficiency of rAAV transduction (Alexander et al., 1994; Kanazawa et al., 2001; Nicolson et al., 2016; Russell et al., 1995). This capability may, perhaps, be an indirect consequence of the cellular reaction to DNA damage: the ‘decoy’ thesis posits that rAAV replication is normally restrained by DNA-damage response proteins, which become preferentially engaged in the work of DNA repair (Choi et al., 2006; Nicolson et al., 2016). Furthermore, doxorubicin may also promote viral nuclear entry through modulation of proteasome function (Yan et al., 2004). We embarked on a series of experiments testing the effect of doxorubicin upon transgene expression mediated by rAAV8 in cat cortex, and observed a remarkable enhancement of transduction following doxorubicin treatment (**Figure 1**, consistent with previous studies utilizing rAAV2 (Zhang et al., 2012; Zhang et al., 2009). Furthermore, doxorubicin enhanced the rapidity of cellular transduction by rAAV: we found that the time required for transgene expression to reach detection level decreased from around 7 days to 3 days, with a sustained increase of expression thereafter (**Figures 1F** and **5**). In short, we infer doxorubicin to have an intracellular action that favors the accrual of stable, dual-stranded AAV DNA as an ‘episome’ within the cell nucleus. This augments expression of the viral DNA, subject to any regulatory elements included within the vector genome. Subsequent investigation served to confirm the generality of this accelerated transduction achievable with doxorubicin.

### The Generality of Doxorubicin-Mediated Transduction Enhancement

AAV has multiple serotypes with varying characteristics and new AAV variants with superior efficacy continue to be developed through vector engineering approaches, that essentially seek to ‘fine-tune’ the binding properties of the capsid to cell surface receptors (Wang et al., 2019). To examine the plurality of doxorubicin enhancement we extended our study of rAAV8 to other AAV serotypes such as rAAV2 and rAAV-PhP.eB. Testing these vectors of different serotypes, we found that all three showed comparable, marked enhancement by doxorubicin, suggesting a serotype-general effect of doxorubicin upon rAAV transduction. For further substantiation, we extended our investigation across species, and found that doxorubicin achieved similar levels of enhancement for transduction by rAAV-PhP.eB in the cortex of macaque monkey, and of the mouse, as it did in the cat (**Figure 4D** and **4E**).

In addition to species- and serotype-generality, we also evaluate whether doxorubicin perturbs the specificity of transgene expression across different cell types, either by interfering with the cell tropism of rAAV vectors, or with the cell-specific regulation of transcription. Our experiments allowed us to compare the action of doxorubicin in conjunction with rAAV vectors carrying either the neuron-specific hSyn or the ubiquitous CAG promoter. Consistent with previous studies (Nathanson et al., 2009; Watakabe et al., 2015), we found a proclivity for transduction that cortical layer 4, characterized by smaller pyramids, showed minimal transduction with rAAV8 under the neuron-specific promoter (hSyn) and the greatest proportional increase with doxorubicin. Most importantly, with hSyn promoter, transgene expression with doxorubicin remained restricted to neurons (**Figure 1D**), giving no evidence of doxorubicin interference with cell-type specificity governed by promoters. With CAG promoter we observed, as expected, additional expression of transgene by astrocytes; doxorubicin treatment induced a similar enhancement in both neurons and astrocytes, but did not alter the negative status of microglia transduction (**Figure S1**). We therefore obtained no indication that doxorubicin can alter the fundamental tropism of an rAAV vectors.

Given what is known of the molecular biology of doxorubicin action (reviewed above), the ensemble of our findings suggests the analogy to chemical catalysis: doxorubicin enhances the speed and level of transgene expression, but does not affect the tropism for cell type (as governed by the capsid), nor alter the specificity of cellular expression (as governed by regulators of transcription). Hence, in principle, doxorubicin enhancement should prove a valuable strategy for optimizing rAAV transduction in many fields of research practice.

### Absence of Appreciable Neurotoxicity at an Effective Clinical Dosage

Cytotoxicity is the foremost concern prohibiting exogenous agents from *in vivo* application to promote rAAV transduction (Nicolson et al., 2016; Russell et al., 1995; Yan et al., 2004). As a chemotherapeutic drug, doxorubicin takes effect in cancer treatment by inhibiting cell proliferation and inducing apoptosis (Tacar et al., 2013). For non-dividing cells, such as neurons, there is less expectation of significant cytotoxicity (although cardiac tissues are known to be affected in high-dose schedules) (Maini et al., 1997). Direct delivery into the brain, rather than intravenous administration, restricts doxorubicin uptake to cells within the local injection area. Our *in vivo* dose-finding study found that doxorubicin significantly enhances rAAV transduction at a doxorubicin concentration of 1∼10 μg/ml (or 1.72∼17.2 μM) (**Figure 2**). At a concentration of 10 μg/ml, we detected little appreciable cytotoxicity: infected neurons maintained healthy morphology and function upon histological examination (**Figures 1D** and **2F**) and *in vivo* imaging (**Figures 5** and **S3**) at 7∼30 days post-injection. Further examinations revealed no apoptosis at the site of injection (**Figure 3A**), nor any inflammatory response (**Figure 3B**). At the behavioral level, our trial animals (mice) also presented normal locomotor activity (**Figure 3D**). This non-neurotoxic concentration of doxorubicin (1.72∼17.2 μM) is within the 10-24 μM range of peak bloodstream plasma concentrations measured in humans following intravenous doxorubicin administration in clinical cancer chemotherapy (Mross et al., 1988). Furthermore, following direct injections into the cortex, or other brain sites, the concentration will fall several-fold upon diffusion through neural tissues. Overall, these results confirm the feasibility of the use of doxorubicin to catalyze rAAV transduction at a dosage effectively matching clinical practice.

### How Would Doxorubicin-Enhanced Transduction Work in Practice?

Our approach mimics many research applications in seeking to transduce neurons at restricted cortical loci (Scanziani and Hausser, 2009). This includes induction of GCaMP6 for two-photon imaging of neural activity, or a marker like GFP to reveal efferent axon terminals, or an effector such as ChR2 to allow optical stimulation of neural activity. Though there have been many successful applications of these methods, enhanced levels of transduction can only be to the advantage of the experimental goals. The demonstration of remote enhancement via retrograde transport upon rAAV2-retro, further expanded potential doxorubicin applications to neural circuit interrogation (Tervo et al., 2016). More importantly, when extended for applications that seek to modulate extensive cortical regions (Eldridge et al., 2016), here the application of doxorubicin could make a strategic difference to the practicality and outcome of a research project, raising the ceiling for what is considered feasible.

Similar considerations arise for subcortical nuclei of large volume such as the putamen. This is particularly the case where the goal is to transduce as much tissue as possible, typically for therapeutic research relating to Parkinson’s disease. In a technical advance over previous work, the most recently completed clinical trial (Christine et al., 2019) employed convection-enhanced delivery (CED) technique, infusing upto 900μl to attain 20-40% coverage of the total putaminal volume. Whilst the assessment of the therapeutic benefits of this treatment was positive, an enhancement of transduction efficiency – as achieved by coadministration of doxorubicin – should give additional benefit.

Finally, systemic delivery via intravenous injection is also generally applied for gene delivery to the nervous system (Foust et al., 2009), and this route provides good safety and convenience, especially for clinical purposes. rAAV9 and several other AAV variants, including the one that we tested in macaque (rAAV-PHP.eB), have been derived specifically for this route to achieve efficient neural transduction within mammalian CNS (Chan et al., 2017; Dayton et al., 2012). However, this route requires substantially high vector load, and faces the challenge of natural immunity (Mingozzi et al., 2007). The enhancement effect of doxorubicin upon transduction would provide great advantages. Combined with BBB opening techniques such as hyperosmotic mannitol or focused ultrasound (Alli et al., 2018; Fu and McCarty, 2016), doxorubicin might prove to be of great avail for rAAV-based gene therapy of neurological diseases. Intravenous co-administration might be applied to address acute conditions such as stroke, exploiting the speedier transgenic expression achievable with doxorubicin. Past gene therapy studies on animal models of stroke have demonstrated benefits from a variety of transgenes, typically administered by direct injection of rAAV, but *prior* to the experimental induction of ischemia (Lu et al., 2012; Sehara et al., 2018). The effective doubling of the rate of transduction achievable with doxorubicin could provide the key to unlock the clinical potential of transgenic therapy in such emergency contingencies.

In summary, our study has demonstrated that application of doxorubicin facilitates rAAV transduction in rodent, carnivore and primate cerebral cortex. This enhancement of doxorubicin on rAAV takes effect at doses well below tolerance without detectable cytotoxicity to neural tissues. Our findings suggest a feasible strategy synergistic with ongoing vector system engineering to optimize rAAV transduction, so to attain more rapid and more efficient transgene expression *in vivo*.

## LIMITATIONS OF THE STUDY

Recombinant AAVs have been widely adopted throughout the biomedical research community, and are utilised by some of the most promising clinical and preclinical trials of gene therapy. Whilst these applications of rAAV have yielded bountiful results, there is scarcely one of them that would not benefit – in terms of the speed, volume or quality of data acquisition – from a simple method of enhancing the efficiency of transgene expression. That is precisely what we describe here, as the main goal of our study was to identify a chemotherapeutic agent – in this case a clinical anti-cancer drug, doxorubicin – for rapidly facilitating rAAV mediated gene expression. There is, plainly, a vast number of potential rAAV vector serotype/promoter/host species combinations and we have only directly investigated a small fraction of this space. Thus, our identification of a ‘catalytic’ action of doxorubicin (that is, free from any interaction with vector serotype, promoter or host species) is presented as a viable (non-falsified) hypothesis as opposed to a verified universal principle. Also, as we have restricted our tests to mammalian cerebral cortex, similar action of doxorubicin in other CNS components (or vertebrate classes) remains a matter of inference. Finally, it is worth repeating that the molecular mechanism by which doxorubicin facilitates rAAV expression was beyond the scope of our study and persists to be unknown; widespread adoption of this technique for multiple applications across several fields of basic and applied biomedical research should incentivize such research, and potentially identify still more potent and/or less toxic agents for enhancing transgene expression.

## RESOURCE AVAILABILITY

### Lead Contact

Information and requests for resources should be addressed to the Lead Contact, Dr. Wei Wang

### Materials Availability

All unique resources generated in this study are available upon reasonable request to lead contact.

### Data and Code Availability

No unique code was generated in this study.

## STAR★METHODS

### KEY RESOURCES TABLE

| REAGANT or RESOURCE | SOURCE | IDENTIFIER |
| --- | --- | --- |
| Bacterial and Virus Strains |  |  |
| rAAV8-hSyn-EGFP | ION Gene Editing Core Facility | N/A |
| rAAV8-CAG-EGFP | ION Gene Editing Core Facility | N/A |
| rAAV-PHP.eB-CAG-EGFP | ION Gene Editing Core Facility | N/A |
| rAAV2-CAG-EGFP | Taitool Bioscience (Shanghai) | N/A |
| rAAV9-hSyn-GCaMP6s | Taitool Bioscience (Shanghai) | Cat# S0528 |
| rAAV2-retro-CAG-tdTomato | Got from Dr. Yuejun Chen' Lab | N/A |
| Chemicals, Peptides, and Recombinant Proteins |  |  |
| Doxorubicin hydrochloride | Sigma-Aldrich | Cat# D1515-10mg |
| Antibodies |  |  |
| Mouse anti-NeuN | Millipore | Cat# MAB377; RRID:AB_2298772 |
| Rabbit anti-GFAP | Invitrogen | Cat# MA119170; RRID:AB_1074604 |
| Rabbit anti-Iba1 | Wako | Cat# 01919741; RRID:AB_839504 |
| goat anti-mouse antibody, Alexa Fluor 555 | Life Technologies | Cat# A21424; RRID:AB_141780 |
| goat anti-rabbit antibody, Alexa Fluor 555 | Life Technologies | Cat# A32732; RRID:AB_2633281 |
| Critical Commercial Assays |  |  |
| One Step TUNEL Apoptosis Assay Kit | Beyotime technology | Cat# C1089 |
| Experimental Models: Organisms/Strains |  |  |
| Mouse: C57BL/6 | SLAC laboratory | N/A |
| Cat | ION animal facility | N/A |
| Rhesus Macaque (Macaca mulatta) | ION Primate Center, Shanghai | N/A |
| Software and Algorithms |  |  |
| ImageJ | NIH | RRID:SCR_003070 |
| GraphPad Prism 7 | GraphPad | RRID:SCR_002798 |
| MATLAB | MathWorks | RRID:SCR_001622 |
| Other |  |  |
| E80i microscopy | Nikon | N/A |
| VS120 virtual slide microscope | Olympus | N/A |
| TiE-A1plus Laser Scanning Confocal microscope | Nikon | N/A |
| Axio Scan Z1 fluorescence microscope | ZEISS | N/A |
| Two-photon microscope | Thorlabs | N/A |
| Axio Scan. Z1 | ZEISS | N/A |
| Ti-Sa laser | Spectra-physics | N/A |

### EXPERIMENTAL MODEL AND SUBJECT DETAILS

#### Animals

All experimental procedures were approved by the Animal Care and Use Committee of the Institute of Neuroscience, Chinese Academy of Sciences. Experimental subjects contained adult cats, C57BL/6 mice and rhesus macaques (Macaca mulatta). Thirty adult cats of both sexes (age: 1-3 years; weight: 2.5-3.5 kg) were obtained through the animal facility of the Institute of Neuroscience. Thirty-three Male C57BL/6 mice of 8 weeks age were purchased from Slac Laboratory Animals (Shanghai). Three macaque monkeys (two males and one female, 6-8 years old, weight: 6.0-9.0 kg) were used in this study, two for cross-species examination and another for neural circuit interrogation. All animals were raised under constant temperature, humidity and automatic light cycles.

#### Virus and drug preparation

The rAAV vectors and sources were the following: rAAV8-hSyn-EGFP, rAAV8-CAG-EGFP, rAAV-PHP.eB-CAG-EGFP (ION Gene Editing Core Facility), rAAV2-CAG-EGFP, rAAV9-hSyn-GCaMP6s (Taitool Bioscience, Shanghai) and rAAV2-retro-CAG-tdTomato (obtained from Dr. Yuejun Chen’ Lab). Doxorubicin (doxorubicin hydrochloride, Sigma-Aldrich) was dissolved in sterile phosphate buffered saline (PBS) and stored at −80°C. For injection, viral solutions were diluted and prepared with PBS just before experiment, which consisted of rAAV and 10 μg/ml doxorubicin (or as indicated). As a control, viral solutions contained only rAAV vector in PBS.

#### Surgery and stereotaxic injection

For virus injection, adult cats were anaesthetized with ketamine hydrochloride (25 mg/kg) and then positioned in a custom-made stereotaxic apparatus with anesthesia maintained by isoflurane (1.5-2.0%). Small skull holes (∼1 mm^2^) were made in both hemispheres at Horsley-Clarke coordinates A5-P7, L2-L3 (**Figure 1A**). Dura incisions were made for introduction of glass pipettes (tip diameter of 40-50 μm) into the cerebral cortex (at a depth of ∼1000 μm from cortical surface). Injections of viral solutions were performed through glass micropipettes at a rate of 90 nL/min by a microinjector (QSI). After injection, micropipettes were kept in place for 10 minutes prior to withdrawal. Pulse oxygen saturation, electrocardiograph, end-tidal carbon dioxide and core body temperature were monitored throughout the surgical procedure. After surgery, animals recovered from the anesthetic and cefotaxime sodium (80 mg/kg) was applied through intraperitoneal injection twice a day in the following seven days.

To examine the effect of doxorubicin on rAAV-mediated neuronal transduction(**Figure 1A**), rAAV8-hSyn-EGFP vector (titer: 1 × 10^12^ vg/ml) was injected into the right and left hemisphere of three cats with or without doxorubicin (10 μg/ml) at a volume of 1000 nl for each injection, respectively. Histological examination was carried out at 30 days post-injection.

To assess the impact of doxorubicin on the kinetics of rAAV transduction (**Figure 1E**), rAAV8-hSyn-EGFP vector (titer: 1 × 10^12^ vg/ml) coupled with doxorubicin (10 μg/ml) or vector alone was administered into the visual cortex of twelve cats at a volume of 1000 nl for each injection, three cats per group. Histological examination was carried out at 3, 7, 15 and 30 days post-injection.

To define the optimal dosage of doxorubicin for neural administration (**Figure 2**), different concentrations of doxorubicin (0.1-100 μg/ml) were injected with rAAV8-hSyn-EGFP (titer: 1 × 10^12^ vg/ml) into cats at a volume of 1000 nl for each injection, with injections of vector only as the visual cortex of six control. Histological examination was carried out at 7 days post-injection.

To assess the cytotoxic effects of doxorubicin upon neural tissues (**Figure 3A-C**), five cats were injected with PBS, doxorubicin (10 μg/ml), and rAAV8 vector (rAAV8-hSyn/CAG-EGFP, titer: 0.5∼1 × 10^12^ vg/ml) with or without 10 μg/ml doxorubicin at a volume of 1000 nl for each injection. Seven days following injection, animals were prepared for TUNEL assay and inflammatory examination.

To determine whether doxorubicin exerts a general agonist effect upon rAAV transduction (**Figure 4A-C**), rAAV-CAG-EGFP vectors (rAAV8, rAAV-PHP.eB and rAAV2, titer: 5 × 10^11^ vg/ml) were injected into the visual cortex of five cats with or without doxorubicin (10 μg/ml) at a volume of 1000 nl for each injection. Histological examination was carried out at 7 days post-injection.

For virus injection in mouse, animals were anaesthetized by 1.5∼2.0% isoflurane or intraperitoneal injection of pentobarbital sodium (40 mg/kg) and mounted in a stereotaxic apparatus with body temperature kept at 37 °C. In the cross-species examination (**Figure 4D,E**), rAAV-PHP.eB-CAG-EGFP vector (titer: 5×10^11^ vg/ml, 1μl) alone or coupled with 10 μg/ml doxorubicin was injected into the primary visual cortex (V1) of four mice (stereotaxic coordinates: 3.1 mm posterior to bregma, 2.5 mm lateral to midline, 0.6 mm vertical from cortical surface) through a skull hole at a rate of 90 nl/min with a glass pipette (tip diameter of 20-30 μm). In the open field test (**Figure 3D-G**), rAAV8-hSyn-EGFP (titer: 1×10^12^ vg/ml, 1μl) alone or coupled with 10 μg/ml doxorubicin was injected into the primary motor cortex (M1) of the right hemisphere of nineteen mice (stereotaxic coordinates: 0.5 mm anterior from bregma, 1.5 mm lateral from the midline, 0.6 mm vertical from cortical surface). For two-photon calcium imaging (**Figure 5 and S3**), a circular craniotomy (∼3 mm diameter) was made to expose the primary visual cortex. rAAV9-hSyn-GCaMP6s (2×10^12^ vg/ml, 1μl) coupled with doxorubicin (10 μg/ml) or vector alone was injected into area V1 of ten mice, at a depth of 0.6 mm beneath the cortical surface. After injection, the dura was removed and a chronic window was made with a custom-made double-layered coverslip covering the cortex. For repeated imaging, a custom-made titanium headplate was implanted with dental acrylic. After surgery, antibiotics were administered by intramuscular injection and animals were given about 7 days (or as indicated) for recovery before experiment.

The three adult macaque monkeys used in this study underwent a similar preparatory procedure. Following induction by ketamine hydrochloride (15 mg/kg), deep anesthesia was maintained with isoflurane (1.5-2.0%) whilst the head of animal was mounted in the stereotaxic apparatus and prepared for craniotomy, durotomy and viral injection. In the cross-species examination (**Figure 4D,E**), rAAV-PHP.eB-CAG-EGFP vectors (titer: 1 × 10^12^ vg/ml) were injected into the primary motor cortex (at a depth of 1500 μm) through glass pipettes (tip diameter of 40-60 μm) with or without doxorubicin (10 μg/ml) at a volume of 2000 nL for each injection. For neural circuit interrogation (**Figure 6** and **S4**), rAAV2-retro-CAG-tdTomato (titer: 4 × 10^13^ vg/ml, 1μl per injection) was injected into the left and right hemisphere of monkey V1 symmetrically, vector alone or coupled with doxorubicin (10 μg/ml). Throughout the surgical procedure, pulse oxygen saturation, electrocardiograph, end-tidal carbon dioxide and core body temperature (37∼38 °C) were monitored. After injection, micropipettes were kept in place for 10 minutes prior to withdrawal and animals were carefully maintained for recovery. Histological examination was carried out at 7 or 42 days post-injection into M1, and V1, respectively.

#### Tissue preparation, histology and image acquisition

Animals were deeply anaesthetized with sodium pentobarbital, followed by transcardial perfusion of 0.9% NaCl and 4% paraformaldehyde (wt/vol) in PBS. Brains were removed and post-fixed with 4% paraformaldehyde in PBS for 12 ∼ 48 hours, and then dehydrated with 30% sucrose (wt/vol) for about 1 ∼ 5 days. For each brain every injection site was identified and marked under a fluorescence stereoscope (Stereo Discovery V20, Zeiss). Fixed brains were cut into 50-μm-thick coronal sections on a freezing microtome. Representative sections from each injection site showing the largest amount of fluorescence were identified and selected for subsequent assay. For immunohistochemistry, sections were first rehydrated with PBS and preincubated with blocking solution (PBS with 1% bovine serum albumin and 0.3% Triton X-100) before incubation with primary antibodies at 4°C overnight. Antibodies against neuronal or glial marker proteins were used as follows: neuron-specific nuclear protein (mouse anti-NeuN, 1:500; Millipore, MAB377), glial fibrillary acidic protein (rabbit anti-GFAP,1:500; Invitrogen, MA119170) and ionized calcium binding adaptor molecule (rabbit anti-Iba1, 1:500; Wako, 01919741). After rinsing with PBS, sections were incubated with secondary antibody (goat anti-mouse antibody, 1:1000, Life technologies, A21424; goat anti-rabbit antibody, 1:1000, Life technologies, A32732) for 2 hours at room temperature. DAPI was added to a final concentration of 1 μg/ml for staining cell nuclei.

Large field images of representative sections from each injection were first examined with Nikon E80i or Olympus VS120 fluorescence microscope using a 10x objective. Confocal fluorescence images were then acquired by Nikon TiE-A1plus confocal laser scanning microscope using a 20x/0.75NA objective. For GFP intensity quantification, the background fluorescence of images was obtained from a remote reference region site and subtracted. GFP signal was identified and calculated from pixels with fluorescence values exceeding the mean background by three standard deviations. To evaluate the effect of doxorubicin, a region of interest (ROI) with 500 × 1300 μm size at the core of each injection was selected for calculation of GFP intensity. For cell number counting, stack images of 10-μm depth were acquired on Nikon TiE-A1plus confocal laser scanning microscope using a 40x/1.3 NA objective. Cells were identified with antibodies identifying cell-type specific marker, NeuN for total neurons, GFAP for astroglia, and Iba1 for microglia. Cell counting was performed manually or automatically using ImageJ (National Institutes of Health).

For neural circuit interrogation in macaque monkey, the brain was prepared for histological procedures according to previously procedures(Markov et al., 2011). Cryoprotection was performed with glycerol gradient perfusions as well as subsequent dehydration with 20% glycerol in PBS. The whole brain was cut into 40-μm-thick coronal sections symmetrically and 1 in 3 sections were taken for imaging and examination. Fluorescence images of three channels (DAPI, EGFP and Cy3) were obtained with ZEISS Axio Scan Z1 fluorescence microscope using a 10x objective. Area parcellation of LGN, V1, V2, V4, V5/MT and other brain regions was performed according to the MRI and histology atlas of rhesus monkey brain(Saleem and Logothetis, 2007), and tdTomato-labelled cells attributed to each area were identified and counted across sections taken every 480 μm manually (**Figure 6 F-H**). For LGN, the sampling interval was reduced to 120 μm and confocal images of 5-μm thickness were taken, with tdTomato-labelled cell bodies of red color counted manually (**Figure 6D**). Pictures were stitched together with Adobe Illustrator software.

#### Two-photon calcium imaging

Mice were anaesthetized by 0.7∼1.0% isoflurane via respiration and fixed with the implanted headplate. Body temperature was maintained at about 37°C with a heating pad, and animal state was monitored using an infrared camera. Images were acquired within an ROI (1024 × 1024 pixels) with a commercial two-photon microscope (Thorlabs) which was controlled by ThorImage software, via a 16× objective (NA = 0.8, Nikon). GCaMP6s-based Ca^2+^ signal was excited with 910-nm wavelength that generated via a mode-locked Ti:sapphire laser (Mai Tai DeepSee, Spectra physics), and collected with filter 525/50. Visual stimuli were generated with the Psychtoolbox for MATLAB and presented monocularly on a 17-inch LCD monitor (Lenove) at a distance of 15 cm in front of the animal. Sinusoidal gratings (contrast = 1; spatial frequency = 0.05 cycle/degree; temporal frequency = 2 cycle/second) were displayed for 3 seconds and drifted at eight directions (0°, 45°, 90°, 135°, 180°, 225°, 270°, 315°) randomly, five repeats per condition. For each condition, images were captured at a frame rate of 7.8 Hz over a period of 8 seconds, including 1 second before stimuli onset. Before experiment, baseline fluorescence was collected for 200 frames without visual stimuli and treated as an indicator of GCaMP6s expression (**Figure 5B**).

Images obtained in our experiment were analyzed with ImageJ (National Institutes of Health) and Matlab (Mathworks). For correcting lateral motion in the imaging data, a rigid-body transformation-based frame-by-frame alignment was applied by using Turboreg plugin (ImageJ software). Cells were identified on the basis of cross-trial variability calculation and manually modified by empirically handed labeling according to the size, shape, and brightness. Fluorescence changes with time of individual cells were extracted by averaging pixel intensity values within each cell in each frame. The fluorescence intensity change (ΔF) evoked by each stimulus (stimulus duration, 3s) was normalized by the pre-stimulus baseline fluorescence (F, 1s). For each stimulus, the mean change in fluorescence (ΔF/F) was calculated by averaging responses to all trials for each stimulus condition. Cells showing significant differences in the fluorescence intensity signals observed during baseline vs. stimulus-presentation periods (P < 0.05, ANOVA) were defined as “responsive cells”. The response strength was determined as the maximal (ΔF/F) across all the stimulus conditions (**Figure S3C**). The orientation selectivity index (OSI) was defined as (1-Var(θ)), where Var(θ) is the circular variance of the responses to the tested directions (**Figure S3D**).

#### TUNEL assay

The TUNEL assay was performed according to the manufacturer’s instructions (Beyotime technology, Shanghai). Briefly, brain sections fixed with 4% paraformaldehyde and permeabilized with 0.3% Triton X-100 were incubated with recombinant termination deoxynucleotidyltransferase (rTdT) for 1 hour at 37°C. The fragmented DNA was labeled at the ends with Cy3-conjugated dUTP to yield red-stained nuclei, indicative of an apoptotic cell. Positive control samples were pretreated with DNase I before TUNEL staining (**Figure 3A**).

#### Behavior test

An ‘open field’ test was conducted according to previously described procedures(Seibenhener and Wooten, 2015). In short, after an 30 minutes acclimation, mice were placed individually in the middle of a 40 × 40 cm open field chamber and allowed to explore freely for 15 minutes (**Figure 3D**). Throughout this period the movement trajectory of the mouse was tracked with a video camera controlled by the EthoVision XT software. Total distance traveled, mean motion velocity, and time in movement were calculated as measures of locomotor activity (**Figure 3E-G**).

### QUANTIFICATION AND STATISTICAL ANALYSIS

Statistical analysis was performed with GraphPad Prism 7 and MATLAB. A test for normality was applied before statistical comparison. If the data conformed to a normal distribution, an unpaired two-tail Student’s t-test or one-way ANOVA followed by post–hoc Tukey Kramer correction were used for comparisons between different groups; if not, the Wilcoxon rank sum test was used. For comparison of transgene expression at three days (**Figure 1F**), both the control group (n = 6) and doxorubicin-treated group (n = 6) had one injection excluded from subsequent analysis upon failure of immunohistochemical detection. For **Figures 3C, 4B, 4C, 4E** and **5B**, data distribution was assumed to be normal and an unpaired, two-tail student’s t-test was used. All data are presented as means ± SEM, and statistical test, n numbers of each group and p values are reported in figure legends. Threshold p values were computed for significance levels: *p < 0.05, **p < 0.01,***p < 0.001.

## ACKNOWLEDGMENTS

This work was supported by the Strategic Priority Research Program of Chinese Academy of Sciences, Grant No. XDB 32060200 to W.W.; Shanghai Municipal Science and Technology Major Project, Grant No. 2018SHZDZX05 to W.W.; National Natural Science Foundation of China Grant No. 31861143032. National Key Research and Development Program of China 2020YFA0112703

## AUTHOR CONTRIBUTIONS

W.W. designed research; HL.G., NN.Y., ZM.S., C.T., performed research; S.S., LL.Q, YL.L, I.M.A., SH.Z, JH.W., and G.Y. contributed analytic tools; HL.G. analyzed data; HL.G., S.S., and W.W. wrote the paper.

## Supplemental Information

**Figure S1.**
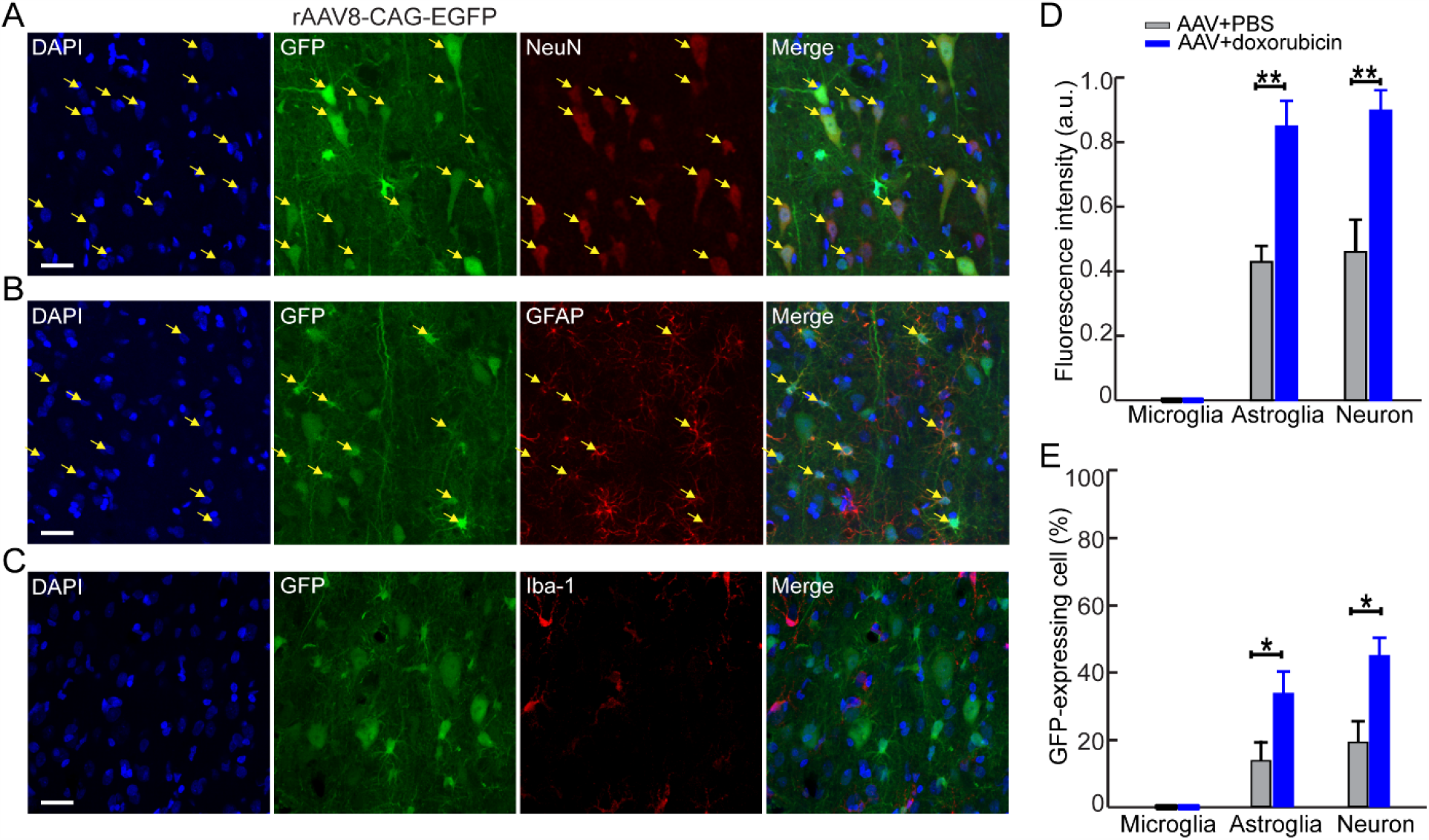
Doxorubicin Promotes rAAV8 Transduction in Neurons with Little Impact on Its Cellular Tropism, Related to Figure 1. (**A-C**) Phenotype of transduced neurons, astroglia and microglia identified with NeuN, GFAP and Iba-1 staining, respectively. Yellow arrows indicate the co-localization of GFP and cell type-specific marker. DAPI, nuclear marker; NeuN, marker of neuron; GFAP, marker of astroglia; Iba1, marker of microglia. Scale bar, 25 μm (**D**) Quantification of GFP intensity in transduced cells in the control and doxorubicin-treated cases. Neuron, p = 0.0061; Astrocyte, p = 0.0068. Unpaired two-tailed t-test. (**E**) Quantification of the percentage of transduced cells in two groups. Neuron, Neuron: 41 ± 4% vs 21 ± 6%; p = 0.038; Astrocyte: 32 ± 6% vs 14 ± 4%; p = 0.033.Unpaired two-tailed t-test. For quantification in **D** and **E**, n = 4 injections from 4 cats, mean ± SEM, *p <0.05,**p <0.01,

**Figure S2.**
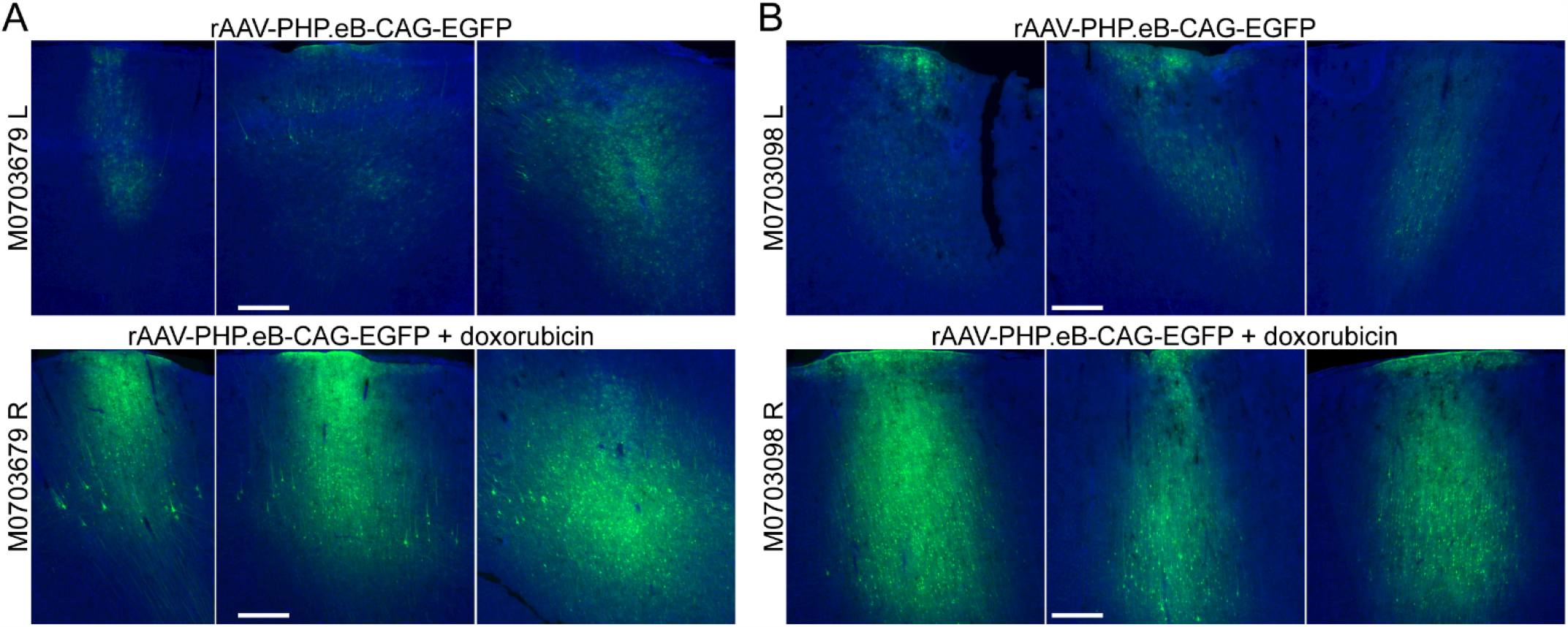
Doxorubicin Enhances rAAV-PHP.eB Transduction in the Motor Cortex of Macaque Monkeys, Related to Figure 4. (**A**) Fluorescence images from three injections of the control and doxorubicin-treated group in M0703679, respectively. rAAV-PHP.eB-CAG-EGFP vector was administered into the left and right motor cortices with or without doxorubicin (10 μg/ml), respectively. GFP intensities were assessed with histological techniques seven days post-injection. (**B**) Similar injections’ results from M0703098. Scale bar, 500 μm.

**Figure S3.**
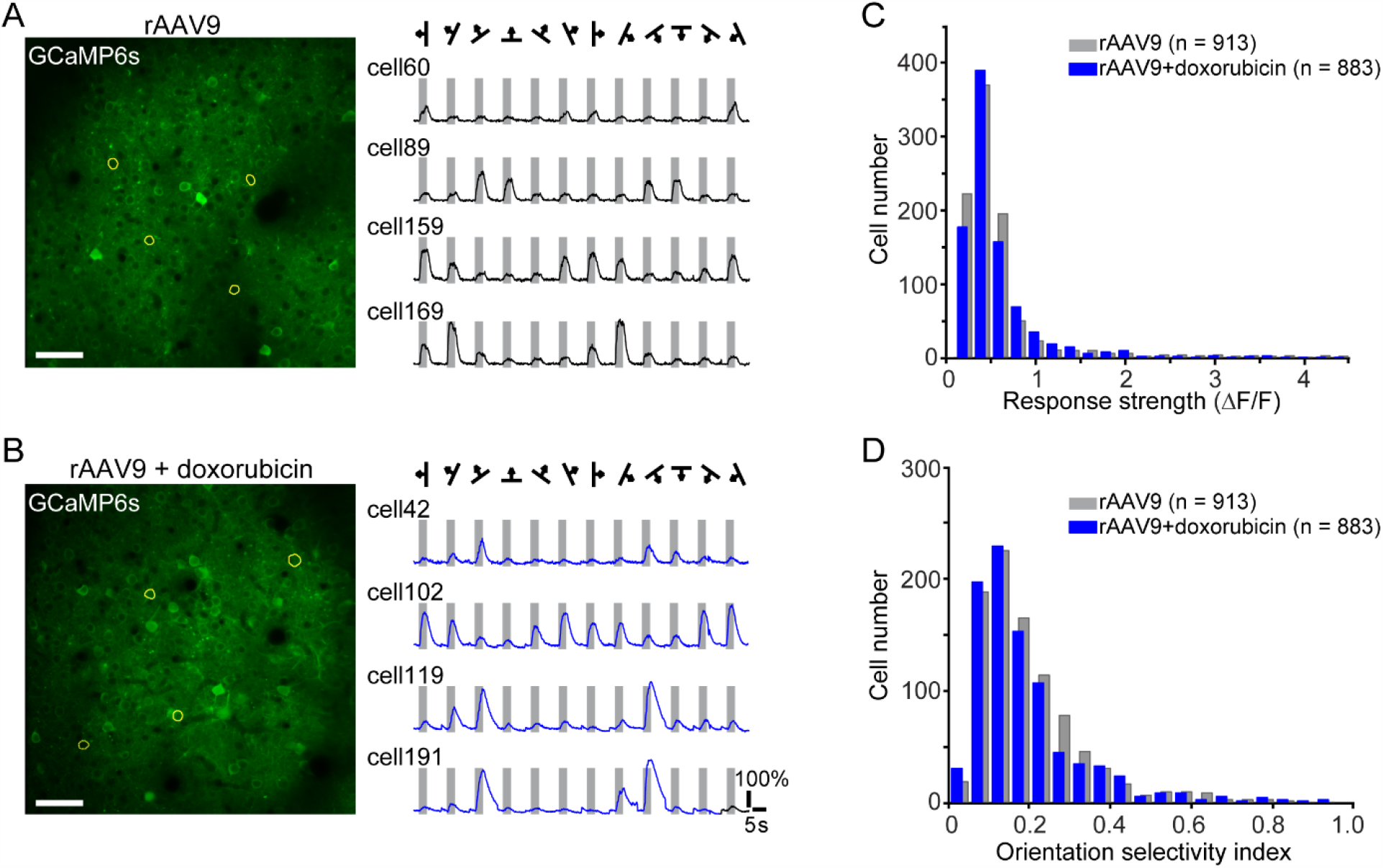
*In Vivo* Two-Photon Calcium Imaging Reveals Little Functional Difference under Experiment Conditions, Related to Figure 5. (**A-B**) *In vivo* two-photon image of Ca^2+^ fluorescence from the control (∼30 day post-injection) and doxorubicin-treated mice (∼15 day post-injection) with comparable expression level of GCaMP6s. Mean fluorescence changes (ΔF/F) of sampled cells (marked by yellow circles) evoked by drifting sine-wave grating stimuli. Scale bar, 50 μm. (**C,D**) Histograms of cell response strength distributions for all responsive cells recorded from the two mouse groups (rAAV9, n = 913 cells; rAAV9-plus-doxorubicn, n = 883 cells). Median of control group 0.43, median of doxorubicin-treated group 0.42; p = 0.25, Wilcoxon rank-sum test. (**E,F**) Histograms of the orientation-selectivity index distributions for all responsive cells recorded from the two groups (rAAV9, n = 913 cells; rAAV9-plus-doxorubicn, n = 883 cells). Median of control group 0.127, median of doxorubicin-treated group 0.116; p = 0.065, Wilcoxon rank-sum test. In **C**-**F**, 5 ROIs per group.

**Figure S4.**
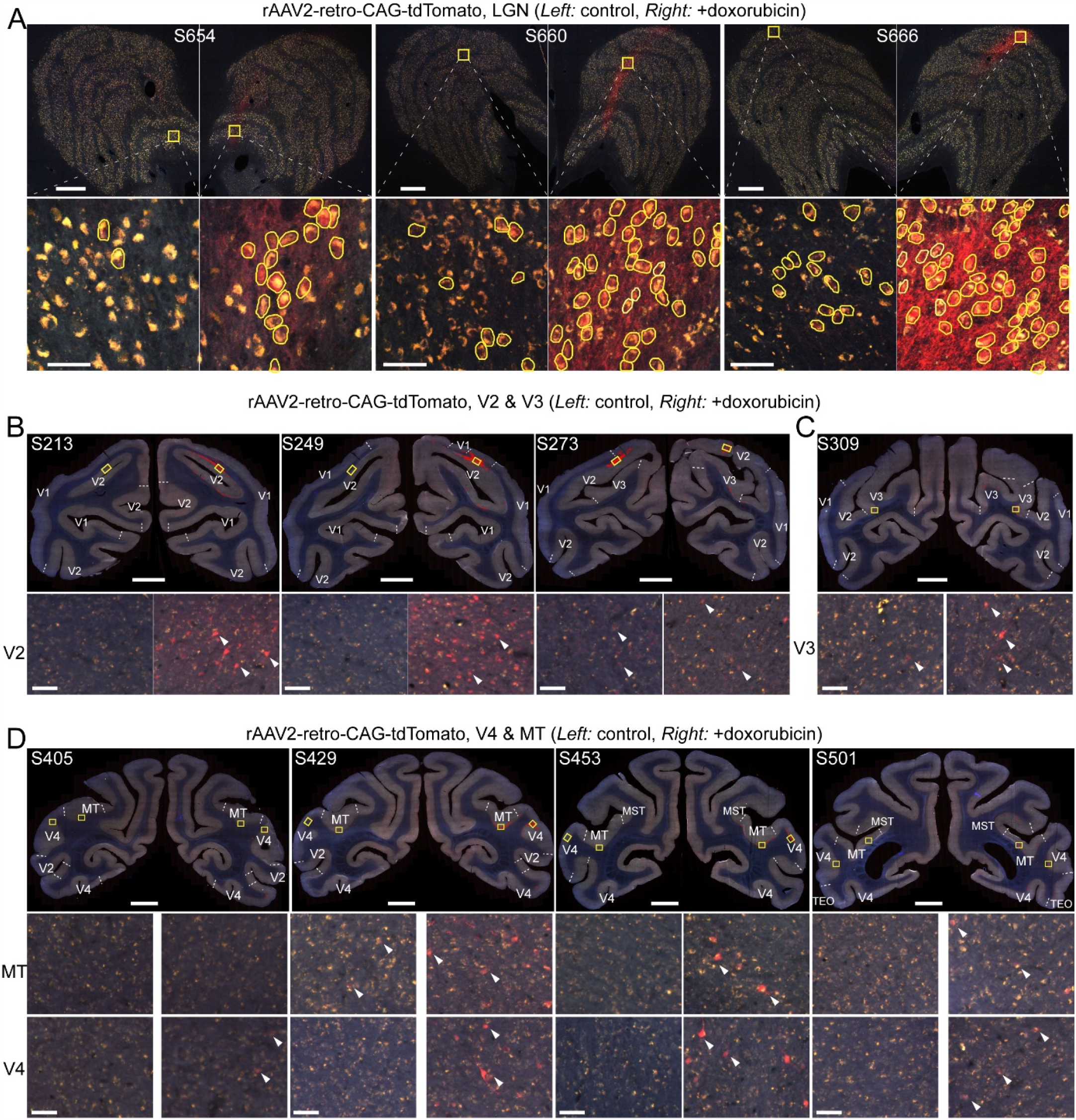
Doxorubicin Enhances rAAV2-Retro-Based Transgene Expression for Neural Circuit Interrogation in the Visual System of Macaque Mo22, Related to Figure 6. (**A**) Schematic representation of LGN sections from hemispheres injected with vector alone (*left*) or vector coupled with doxorubicin (*right*). Upper panel: whole coronal image of LGN in different sections (S654, S660 and S666); scale bar, 500 μm. Lower panel: transduced neurons from the above ROIs are marked with white circles; scale bar, 50 μm. (**B**) Schematic representation of V2 sections from the left and right hemisphere. (**C**) Schematic representation of V3 from the left and right hemisphere. **(D)** Representative images of V4 and MT from the left and right hemisphere. Upper panels in **B, C** and **D** show whole coronal images with transduced cells marked by red dots; scale bar, 5000 μm. Lower panels (middle and bottom panels in **D**) show transduced neurons from ROIs (boxed in the upper panel) of V2, V3, V4 and MT marked by white arrowheads; scale bar, 50 μm.

